# Effect of mannose modified chitosan on uptake of nanoparticles by macrophages

**DOI:** 10.1101/620906

**Authors:** David Lowsberg, John Smith

**Author notes:** Address: 631 Sumter Street, Columbia, SC 29208.

## Abstract

This report provided a new method to prepare and evaluate mannose-modified chitosan-coated lactic acid-glycolic acid copolymer (PLGA) nanoparticles, and to investigate their effects on macrophage toxicity and macrophage uptake. The PLGA nanoparticles loaded with ovalbumin (OVA) were prepared by double emulsion method. The size and zeta potential of the nanoparticles were determined by laser granulometry after mannose-modified chitosan coating. The nanoparticles were observed by transmission electron microscopy. The appearance of the form, the BCA method to determine the OVA content, calculate the drug loading and release. The OVA nanoparticles labeled with fluorescein isothiocyanate (FITC) were co-incubated with macrophages (RAW 264. 7), cell viability was determined by MTT assay, and uptake was examined by fluorescence microscopy. Results The size and ζ potential of OVA-PLGA nanoparticles increased with the increase of chitosan coating concentration (P < 0.05), and OVA drug loading range was 7. 2% to 8. 4%. Chitosan and mannose modified chitosan coating FITC-OVAPLGA nanoparticles and RAW 264. 7 After incubation, there was little effect on cell viability (P > 0.05), but it significantly promoted macrophage uptake by FITC-OVA-PLGA nanoparticles (P < 0.05).

## Introduction

Macrophages and dendritic cells are specialized antigen-presenting cells (APCs) in the body and play an important role in the prevention and treatment of natural and acquired immunity, inflammation and tumors [1–3]. By increasing the targeting and accessibility of vaccines to APCs^1-6^, the goal of promoting immune response and reducing side effects can be achieved [4, 5]. Macrophages have the property of phagocytizing foreign particulate matter, and if a passive targeting strategy such as receptor-mediated nanocarrier administration is employed, the targeted delivery of antigen can be further improved^7-20^. It has been reported that a large number of C-type lectin family transmembrane protein-mannose receptors are expressed on the surface of macrophages to recognize polysaccharide molecules and soluble glycoproteins on the surface of cells or pathogens, and through ligand-receptor binding, Endocytosis or phagocytosis eliminates pathogens and excess glycoproteins in the body [6]. Yeeprae et al [7] proposed the effect of ligand density on cell recognition and phagocytosis by preparing mannose-modified O/W emulsion. Zhou et al [8] applied mannose modified chitosan microspheres to improve the immune response of hepatitis B virus DNA, indicating that mannose modification improved the release of chitosan microspheres and membrane permeability. In this study, ovalbumin (OVA) was used as a model antigen to prepare biodegradable polylactic-coglycolic acid (PLGA) nanoparticles by double emulsion method, and chitosan was modified by self-made mannose. After the coating treatment, the effect on the uptake function of macrophages was investigated to provide a basis for the research of new vaccine preparations.

## Materials and Method

Reagent fluorescein isothiocyanate (FITC)-OVA and mannose modified chitosan (Man-CS) made by our laboratory; lactic acid-glycolic acid copolymer (PLGA, 50: 50, molecular weight 20 000) was purchased from Jinan Biotech Co., Ltd.; Chitosan (CS, molecular weight 8 000 ∼ 10 000) purchased from Nantong Xingcheng Biological Products Factory; 2,2’-biquinoline-4, 4’-dicarboxylic acid disodium (BCA, purity: BR) purchased from Suzhou Industrial Park Yake Chemical Reagent Co., Ltd.; DMEM high sugar medium purchased from Gibco; Newborn bovine serum purchased from Hangzhou Sijiqing Biotechnology Co., Ltd.; Trypsin, glutamine (purity: BR) purchased from Sinopharm Chemical Reagent Co., Ltd.; Hepes and DAPI staining solution were purchased from Biyuntian Biotechnology Research Institute, Haimen; Macrophage (RAW 264. 7) was purchased from Shanghai Library of Chinese Academy of Sciences. Preparation of Man-CS coated PLGA nanoparticles Preparation of OVA-PLGA nanoparticles Reference [9]: Add 250 μl of 10 mg OVA in polyvinyl alcohol (PVA) solution (mass concentration 3%) to PLGA acetic acid In the ethyl ester solution, the colostrum (W1 /O) was prepared by ultrasonic bathing for 10 s (200 W, working 1 s / interval 1 s). Then add colostrum to the PVA solution (mass concentration 3%), stir evenly in the ice bath (15 000 r / min × 5 min), phacoemulsification (400W, work 1 s / interval 1 s), forming oil-in-water Water-containing double emulsion (W1 / O / W2). Finally, the double emulsion was transferred to a PVA solution (20 ml, mass concentration of 0.5%), and magnetically stirred to a completely volatile organic solvent at room temperature, centrifuged (3 000 r/min × 10 min), and the precipitate was discarded. The supernatant was centrifuged (19 000 r/min × 20 min), and the precipitate was collected and washed 3 times with deionized water. Pre-frozen (−70 ° C), freeze-dried, that is, PLGA nanoparticles. CS or Man-CS coating: The above double emulsion is directly added to a PVA acetic acid solution containing a certain concentration of CS or Man-CS (mass concentration of 0.5%), the organic solvent is volatilized at room temperature, centrifuged, washed 3 times^21-27^, frozen Dry is available. Nanoparticle preparation evaluation. Nanoparticle size and ζ Potential Determination The nanoparticle colloidal solution or lyophilized nanoparticles dispersed in distilled water were taken, and the size of the nanoparticles and the zeta potential were measured by a laser particle size analyzer. Nanoparticle morphology observation The nanoparticle solution was diluted to the appropriate concentration, stained with 2% phosphotungstic acid, dropped onto a copper mesh, and observed under a transmission electron microscope. Encapsulation rate and drug loading determination Precision weighed lyophilized nanoparticles 20 mg, added to 1 ml 5% sodium lauryl sulfate / 0. In a 1 mol / L NaOH solution, shake at a constant temperature of 25 ° C for 4 h, centrifuge (10 000 r / min) for 10 min. The OVA content in the supernatant was determined by the BCA method, and the loading amount (LA) and the entrapment efficiency (EE) were calculated according to the following formula: LA = the amount of protein in the nanoparticles. The mass of the nanoparticles × 100% EE = amount of protein in the nanoparticles. Total amount of protein added × 100%. Nanoparticle release in vitro Precision weighed lyophilized nanoparticles 50 mg Dispered in 1. In a 5 ml EP tube, add 1 ml of poloxamer 188 solution (mass concentration 0.02%, PBS, pH = 7.4), shake at 37 °C, and accurately remove at a predetermined time. 5 ml, centrifuge (19 000 r/min) for 15 min. The supernatant was taken, the protein concentration was determined by the BCA method, the release was calculated, and the same volume of the release medium was added. Man-CS-OVA-PLGA Nanoparticle Cytotoxicity Test Rapidly thaw frozen macrophages (RAW 264. 7) at 37 °C, add 7 ml of DMEM high glucose medium, centrifuge twice at 1 000 r/min. Add DMEM high glucose medium containing 10% calf serum (containing 100 U / ml penicillin and streptomycin) in 37 ° C, 5% CO 2 incubator, change the culture medium on the 2nd day, the cells are covered 70% ∼ 80 When %, use 0. 25% trypsin digestion. The cells were seeded in 96-well plates at 5 × 103 /well^28-32^, and cultured until the cells were attached. The experimental group was added with free FITC-OVA, PLGA nanoparticles containing FITC-OVA, and chitosan-coated PLGA (CS-PLGA). Nanoparticles and mannose-modified CS-coated PLGA (Man-CS-PLGA) nanoparticles. Another cell blank control well is set, and 6 replicate wells are provided for each well. After culturing for a certain period of time in the incubator, 96-well plates were taken, 10 μl of medium was aspirated per well, and 10 μl of tetramethyl azozolium salt (MTT) was added, and incubation was continued for 4 h. All the supernatant was aspirated, and 100 μl of SDS-HCl was added to each well overnight. The absorbance (A) was measured at 570 nm with a microplate reader, and the cell viability was calculated by adjusting the blank of the medium to zero. 5 Macrophage uptake experiments. The trypsin-digested logarithmic growth macrophages were seeded in 6-well plates (5 × 105 /ml) at 2 ml per well and cultured overnight. Then, PBS, free FITC-OVA, FITC-OVA-PLGA nanoparticles, CS-PLGA nanoparticles and Man-CS-PLGA nanoparticles were added and incubated for 8 h. The supernatant was discarded and washed 3 times with PBS and observed under a fluorescent microscope. In the above test, the concentration of FITC-OVA was 20 μg / ml. Statistical processing. The measurement data is expressed as x ± s, using SPSS 16. 0 Statistical software for one-way ANOVA, P < 0. 05 indicates that the difference is statistically significant.

## Results and discussion

The effect of CS or Man-CS coating on PLGA nanoparticle size, ζ potential and drug loading is shown in Table 1. It can be seen from Table 1 that the chitosan coating increases the particle size of PLGA nanoparticles and tends to increase with the increase of chitosan concentration (P < 0.05). The zeta potential of the PLGA nanoparticles was negative and was converted to a positive value after chitosan coating and was affected by the chitosan concentration (P < 0.05). However, the chitosan coating concentration had little effect on the drug loading of OVA (P > 0.05).

**Figure 1.**
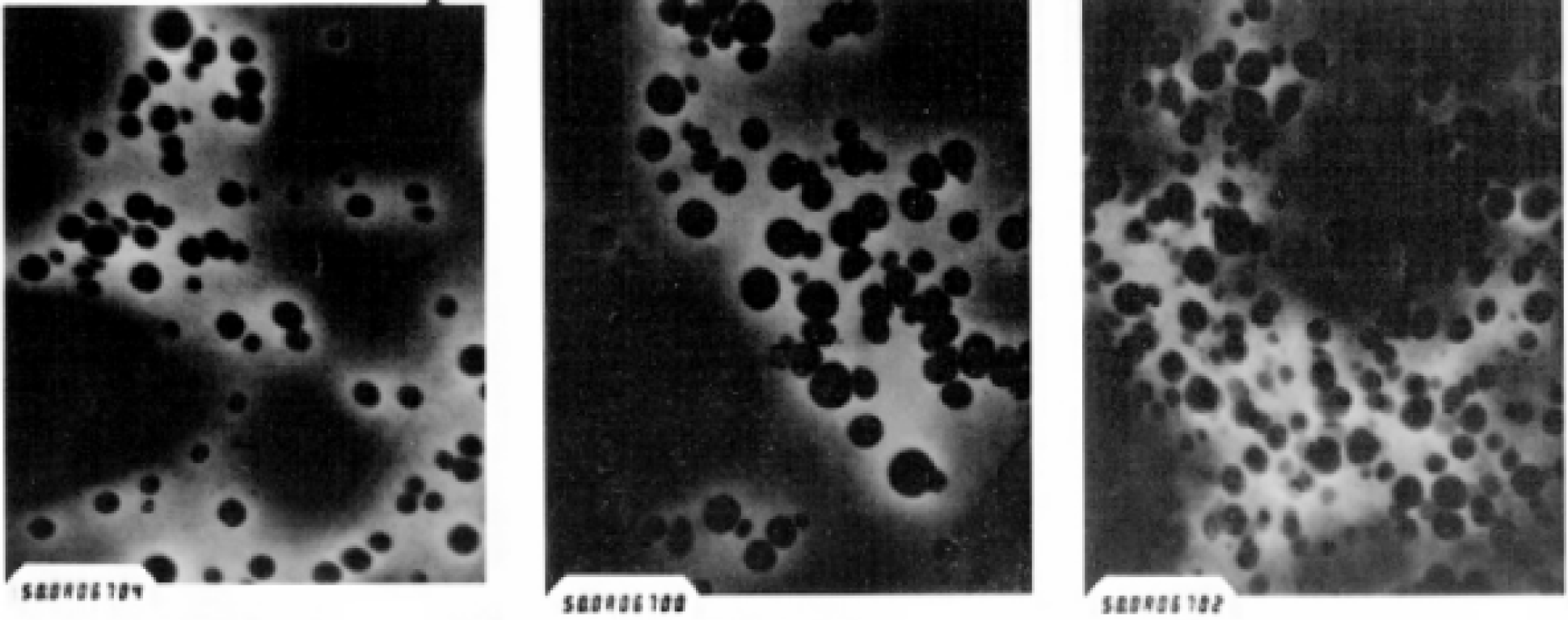
TEM analysis of nanoparticles.

The release of OVA-PLGA nanoparticles showed significant burst and sustained release characteristics. The 1 h release rate of OVA exceeded 30%, followed by release near zero-order velocity, and the release rate reached 80% at 30 d. OVA-PLGA via CS After coating with Man-CS, OVA burst release was reduced, but the percentage of total OVA release was also significantly reduced (both P < 0.05) (Figure 2).

**Figure 2.**
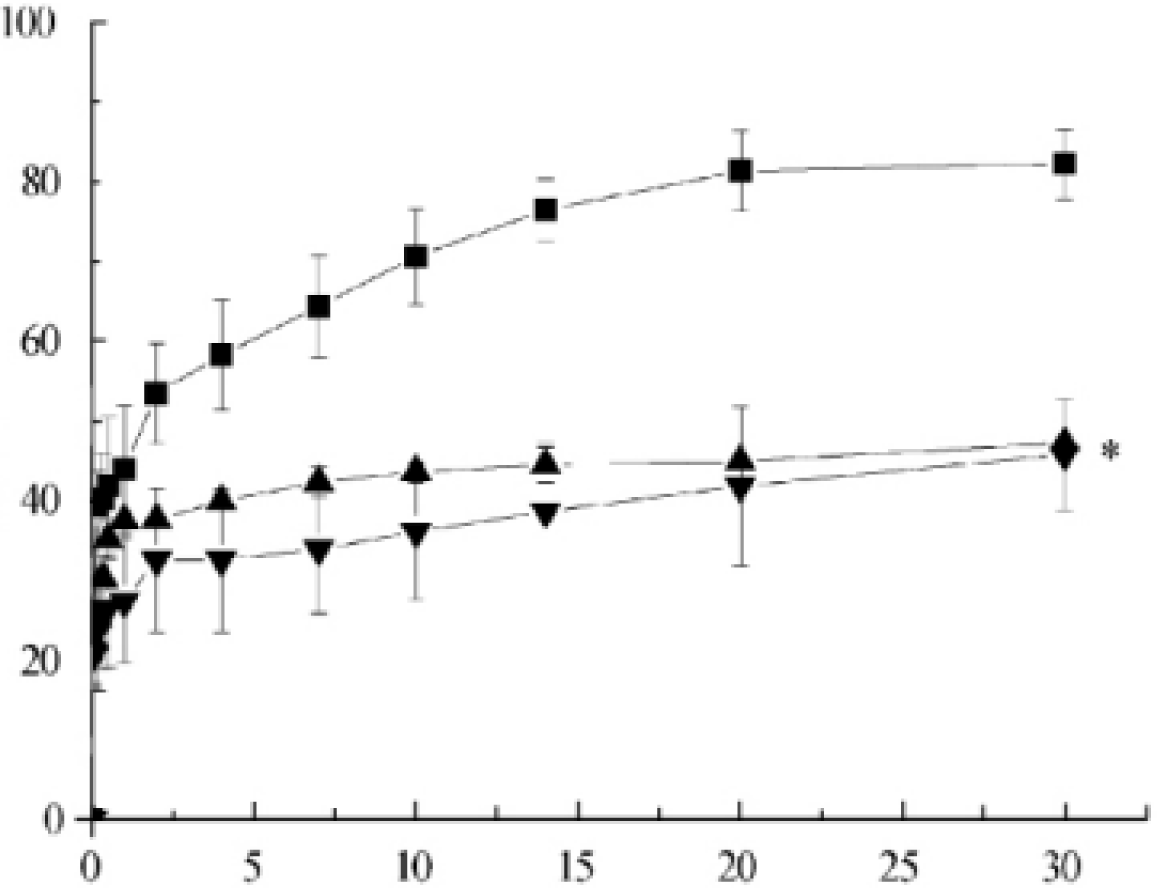
Releasing kinetics of nanoparticles.

Macrophages were incubated with free FITC-OVA, FITCOVA-PLGA nanoparticles, CS-coated PLGA (CS-PLGA) nanoparticles, and mannose-modified chitosan-coated PLGA (Man-CS-PLGA) nanoparticles. At 8 h, the viability of the cells measured by the MTT method is shown in Figure 3. As can be seen from Figure 3, free FITC-OVA, FITC-OVA-PLGA nanoparticles, CS-PLGA nanoparticles, and Man-CS coated PLGA nanoparticles at a concentration of 10 to 80 μg / ml were co-incubated with macrophages. Except for Man-CS-PLGA nanoparticles with 80 μg / ml concentration, cell viability was higher than 90%. Analysis of variance showed no significant difference (P > 0.05).

**Figure 3.**
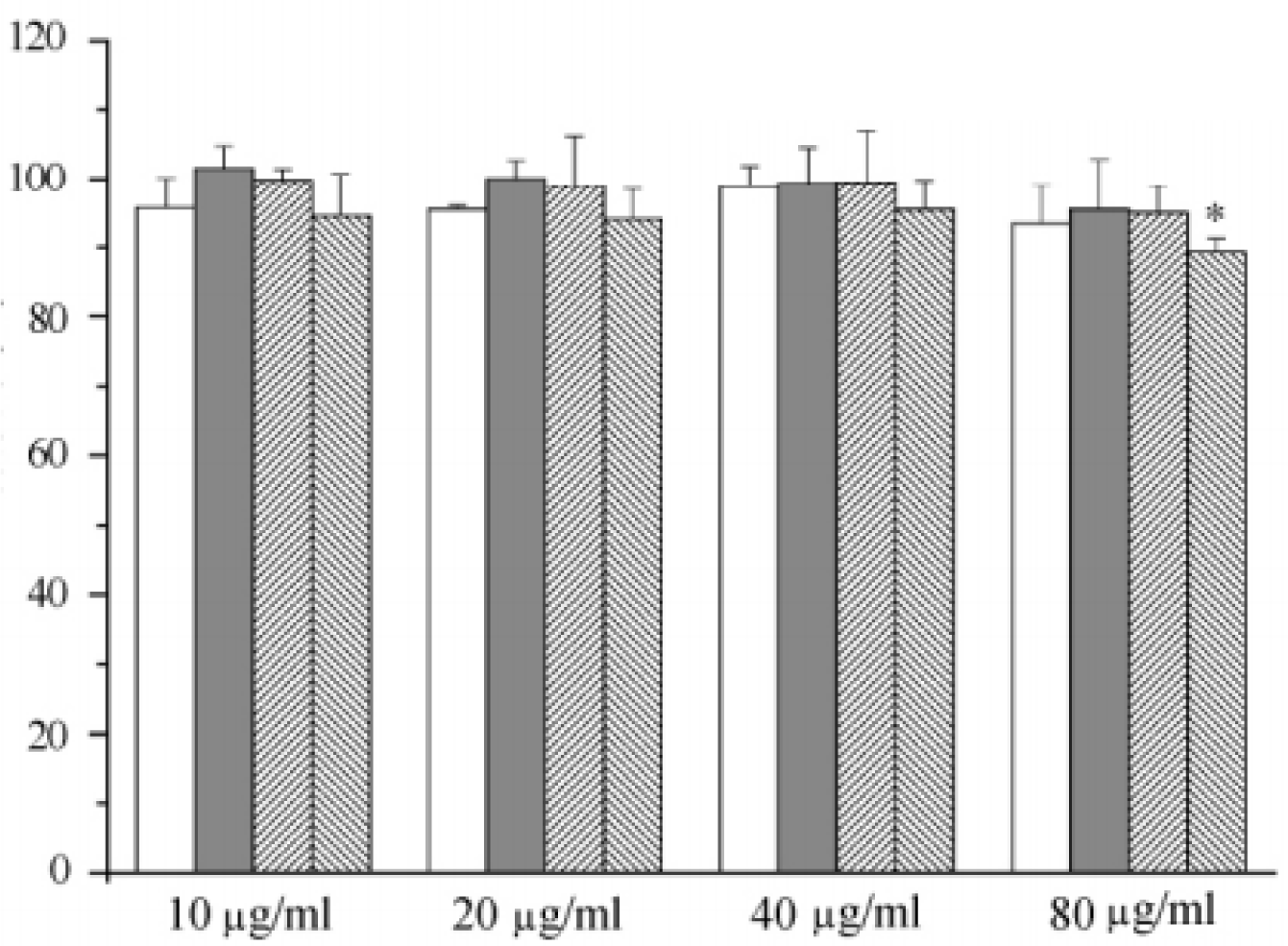
Viability analysis of different nanoparticle formulations at different concentrations.

The effect of Man-CS coating on the uptake function of macrophage FITC-OVA-PLGA nanoparticles is shown in Figure 4. As can be seen from Figure 4, the fluorescence intensity of macrophages was weak after co-incubation with free FITC-OVA solution. However, the fluorescence intensity of macrophages was significantly increased after incubation with FITC-OVA-PLGA nanoparticles. The macrophages co-incubated with Man-CS coated FITC-OVA-PLGA nanoparticles showed the strongest fluorescence, followed by CS coated FITC-OVA-PLGA nanoparticles.

**Figure 4.**
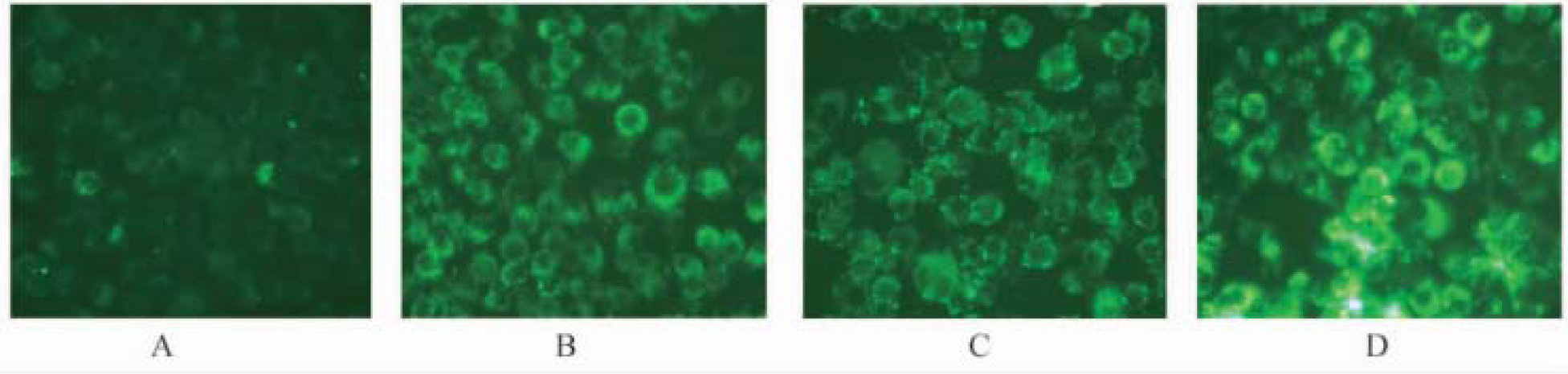
Cell uptake analysis by fluorescent microscopy

In this study, CS and Man-CS coated FITCOVA-containing PLGA nanoparticles were prepared and their formulation properties were evaluated. The particle size of the nanoparticles is (256 ± 30) ∼ (305 ± 16) nm, and the drug loading is (7. 2 ± 1. 5) % ∼ 8. 4 ± 0. 6) %. After the coating treatment, the size of PLGA nanoparticles increased, and the surface charge of the particles also changed from negative to positive, and was affected by the concentration of CS. The coating treatment reduced the burst effect of OVA, but also hindered the subsequent release process, indicating that its formulation and preparation process has yet to be optimized.

MTT assays after co-incubation of macrophages with different nanocarriers showed that cell viability was above 90% with no significant cytotoxicity. The FITC-OVA uptake rate of macrophages from low to high was followed by FITC-OVA solution < FITC-OVA-PLGA nanoparticles < CS-FITC-OVA-PLGA nanoparticles < Man-CSFITC-OVA-PLGA nanoparticles. The uptake of macrophages by nanoparticles is higher than that of solutions, which is related to the different mechanisms of uptake of liquids and nanoparticles [10]. In addition, since the surface of the cell membrane is generally negatively charged, the CS coating causes the surface of the FITC-OVA-PLGA nanoparticle to be positively charged, and then the expression of the receptor-associated mannose ligand on the surface of the macrophage significantly promotes cell uptake.

## Conclusion

The mannose-modified cationic nanoparticle system carrying the model antigen was preliminarily established, which provided a basis for the study of anti-targeting cell targeting in vivo. Future efforts will focus on transferring this promising technology to the clinical settings

